# Anuran call properties as reliable indicators of environmental suitability for reproduction

**DOI:** 10.1101/2024.06.17.599371

**Authors:** Julianne E. Pekny, Brian D. Todd, Eric Post

## Abstract

The annual onset of breeding activity by animals in seasonal environments is often accompanied by auditory signals such as stridulations in insects and vocalizations in birds, mammals, and anurans. In ectotherms, the seasonal timing of such activity by males signals to females the general favorability of environmental conditions for reproduction. Complementary to this, the acoustic characteristics of male auditory signals are presumed to indicate, primarily, the status or quality of males as mates or territorial competitors. Here, using male anurans as a case study, we present the novel hypothesis that characteristics of auditory signals that are modulated by temperature may also serve as bioclimatic indicators of the suitability of proximal abiotic conditions for reproduction by females. According to this hypothesis, thermal constraints on characteristics such as call rate and duration that are insuperable by males may imbue their calls with features that facilitate tracking of environmental conditions by females independently of information intended for communication by males. We integrate findings from empirical studies spanning multiple fields of ecology and evolution to demonstrate how temporal variation in call properties may directly influence female reproductive behavior and physiology. Finally, we outline how this proposed mechanism may enable environmental tracking, provide guidelines for future research to experimentally test this hypothesis, and discuss how findings from this research can translate into actionable conservation management for species of concern.

## INTRODUCTION

Auditory cues like birdsong and the chorusing of frogs are often synonymous with the changing seasons. To conspecifics, these signals indicate the presence and quality of potential mates or competitors. Interspecific variation in the properties of acoustic signals facilitate species recognition (Ryan & Rand, 1993), while intraspecific variation functions in mate choice and (Searcy & Andersson, 1986; Gerhardt, 1991) and male competition (Arak, 1983; Ramer et al., 1983). Information about a signaler’s environment may also be encoded in these cues, however, broadcasting not only the signaler’s quality as a potential mate or competitor, but also the suitability of conditions at the signaler’s location for reproduction. Thus, in seasonal environments or spatially heterogeneous environments, auditory cues may facilitate environmental tracking and mediate spatiotemporal patterns of reproduction.

Sound production, like many other biological processes, is physiologically limited by body temperature, restricting the duration of acoustic signals and the rate at which they can be produced (Gillooly & Ophir, 2010). Because body temperature in ectotherms is constrained by environmental temperature, the rate and length of acoustic signals emitted by ectotherms are influenced by the signaler’s immediate environment. Therefore, because temperature is a constraining factor for ectotherm reproduction and directly influences the properties of acoustic signals, these signals communicate information about the suitability of a caller’s environment for reproduction. As reproduction is energetically costly (Wootton, 1985; Audzijonyte & Richards, 2018), and often risky (Magnhagen 1991), information about the condition of breeding sites prior to arrival may enable receivers to optimize reproductive success. In addition to selecting among mates, individuals must choose which breeding microhabitat to approach across a heterogeneous landscape or, in seasonal environments, when to time their arrival to coincide with optimal abiotic conditions. Cues that signal environmental conditions at these microhabitats may thus influence reproductive patterns across both space and time.

The timing of amphibian reproduction in seasonal environments is dynamic and amphibians exhibit the greatest documented rates of phenological advance in relation to climate change yet reported among vertebrates (Parmesan, 2007; Todd et al., 2011), suggesting that proximate environmental conditions cue amphibian reproduction. Temperature and rainfall are correlated with reproduction in many amphibian systems (Todd et al., 2011; While & Uller, 2014; Ficetola & Maiorano, 2016), but the organism-level mechanisms contributing to this relationship remain unresolved. Recent studies of anurans—the frogs and toads—show that social factors such as male chorusing may be better indicators of female anuran reproductive behavior and annual phenology than abiotic environmental conditions (Höbel, 2017; O’Brien et al., 2021). Using anurans as a case study, we highlight a previously unexplored possible cue for reproduction that may be broadly applicable to terrestrial ectotherms like anurans, cicadas, and other insects that rely on acoustic signals for breeding: environmentally mediated variation in the gross temporal properties of conspecific acoustic signals.

### Box 1

**Environmentally mediated variation in anuran acoustic signal properties**

One type of anuran vocalization, advertisement calls (*sensu* Wells, 1977), are emitted by males at breeding sites and used by females to find and select mates. Anuran advertisement calls are emitted at regular intervals and may contain multiple notes, which can be further divided into bursts of sound energy called pulses (Figure 1). The four most widely studied properties of these calls are dominant frequency, call rate, call duration, and pulse rate. Among these, dominant frequency is a spectral property that is typically static among individuals, whereas call rate and duration, gross temporal properties, are often dynamic (Köhler et al., 2017). Temporal properties are strongly correlated with the caller’s immediate environment—while male call rate and/or duration can be behaviorally modified over brief periods when in the presence of competitors, these properties are physiologically limited by muscle contraction, which in anurans is constrained by ambient temperature. Correspondingly, the temporal properties of anuran advertisement calls are, in general, significantly correlated with ambient temperature while spectral properties show no, or fewer, significant relationships with temperature (reviewed in Köhler et al., 2017). Based on the dynamic, intraspecific variation of gross temporal properties and their strong, repeatable correlation with ambient temperature, we focus on the effects of gross temporal properties throughout this article but acknowledge that spectral and fine scale temporal properties may also communicate information about abiotic conditions at breeding sites in some systems.

**Figure 1.**
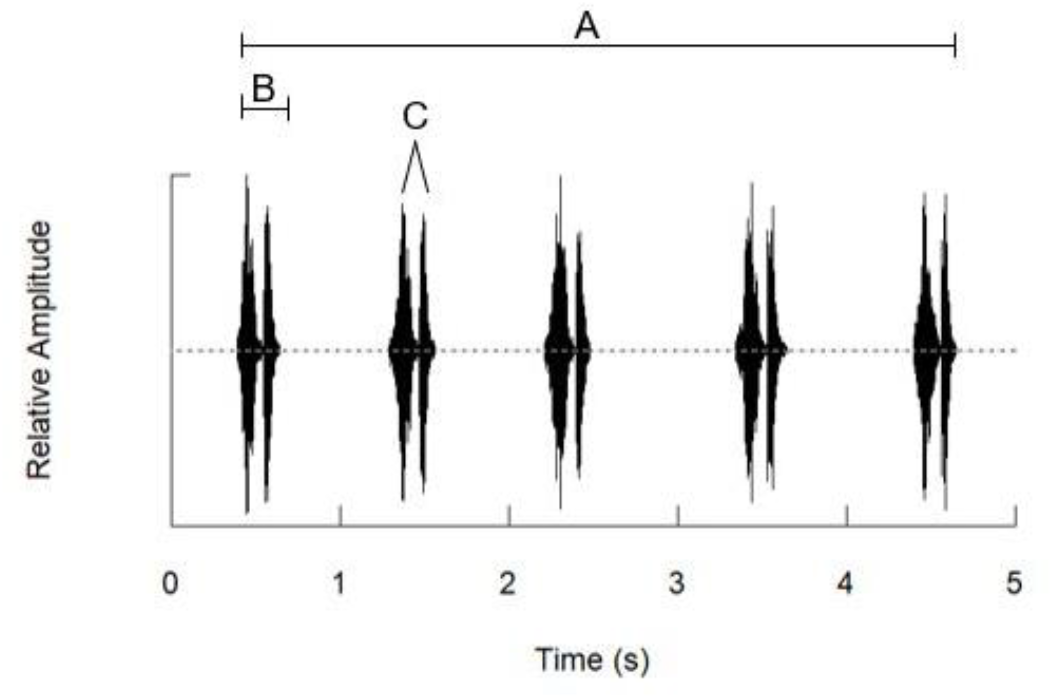
Oscillogram of a male Sierran chorus frog (*Pseudacris sierra*) advertisement call series, recorded in Napa County, California, USA, March 2022. A call series (A) consists of consecutive calls (B) emitted at regular intervals. Calls can contain multiple notes (C), which are created from bursts of sound energy, called pulses.

### Box 2

**Anuran advertisement call properties as reliable indicators of call site environmental conditions: an illustrative example from our research**

To determine how variation in the gross temporal properties of male advertisement calls may indicate the suitability of a breeding site for reproduction, we conducted a field experiment with male Sierran chorus frogs (*Pseudacris sierra; N=35*) in which individuals’ calls were recorded from three different experimentally controlled water temperature treatments (methodological details and further results available in the online supplementary information). The treatments in this experiment correspond to water temperatures above and below 8.0°C because this is the lowest temperature in which pairs of Pacific chorus frogs (*P. regilla*), the closest congener of *P. sierra* (see Recuero et al., 2006; Barrow et al., 2014; Bonett et al., 2017), have been observed in amplexus (Cunningham & Mullally, 1956). Each frog was tested in each water temperature treatment following an acclimation period of at least 24 hours and males were in contact with the water in all trials. The gross temporal properties of calls recorded during behavioral trials were measured using RavenPro software. We found that both call rate and call duration were significantly affected by water temperature (Figure 2), with an increase in call rate and decrease in call duration as water temperatures increase (see supplementary materials for model structure and parameter estimates). Conversely, no significant relationships among gross temporal properties and any of the measured indicators of male quality, including size, mass, and condition were detected. Thus, the observed variation in call rate and call duration appears to be a reliable indicator of water temperature at calling sites in this system. Consequently, female *P. sierra* are exposed to conspecific advertisement calls that vary significantly in gross temporal properties when emitted from breeding sites that vary in water temperature, even by the same male (Figure 2).

**Figure 2.**
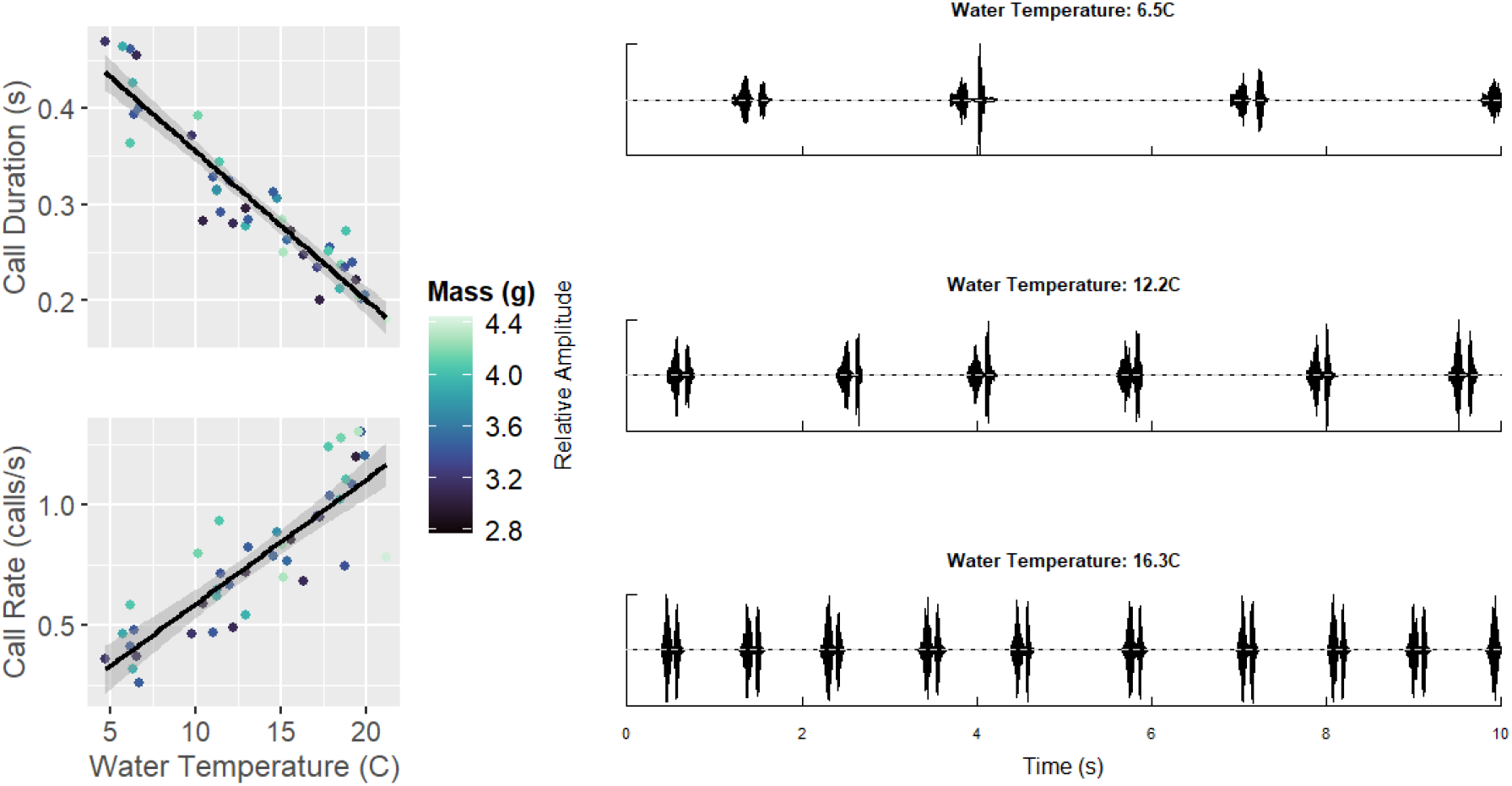
The relationships among water temperature and two gross temporal properties of Sierran tree frog (*Pseudacris sierra*) advertisement calls, call rate and call duration, in a field experiment with wild-caught males at Quail Ridge Ecological Reserve (Napa County, California, USA). Water temperature showed a significant positive relationship with call rate and significant negative relationship with call duration (left), with no detectable effects of any measured index of male quality, including mass, length, or condition (the ratio of mass to length). Thus, advertisement calls recorded in this experiment, emitted from the same male but in different water temperatures, showed variation in call rate and call duration (right).

### GROSS TEMPORAL PROPERTIES OF ADVERTISEMENT CALLS AS BIOCLIMATIC INDICATORS OF BREEDING SITE SUITABILITY

We hypothesize that variation in the gross temporal properties of advertisement calls (Box 1) may act as a bioindicator of abiotic conditions at breeding sites, providing a cue for environmental tracking through two potential mechanisms: stimulation of female reproductive physiology and behavior (Figure 3; we review known effects of acoustic signals on female anuran physiology below in “Female response to advertisement calls with different gross temporal properties”). As gross temporal properties are limited by abiotic environmental conditions, advertisement calls emitted by males at the breeding site provide females with information about the suitability of a caller’s environment for reproduction in addition to the presence and quality of potential mates. In temperate environments where seasonality is characterized by thermal conditions, advertisement calls provide temporally-coherent and consistently updated information each night as seasonality progresses. For example, as the water in snowmelt-fed mountain lakes across the western United States warms as spring unfolds, advertisement calls emitted by male Pacific chorus frogs (*Pseudacris regilla*)—a common anuran which typically calls when in contact with water (Vélez & Guajardo, 2021)—from within those lakes will generally show increasing call rates, providing cotemporary information on current water temperature to females daily.

**Figure 3.**
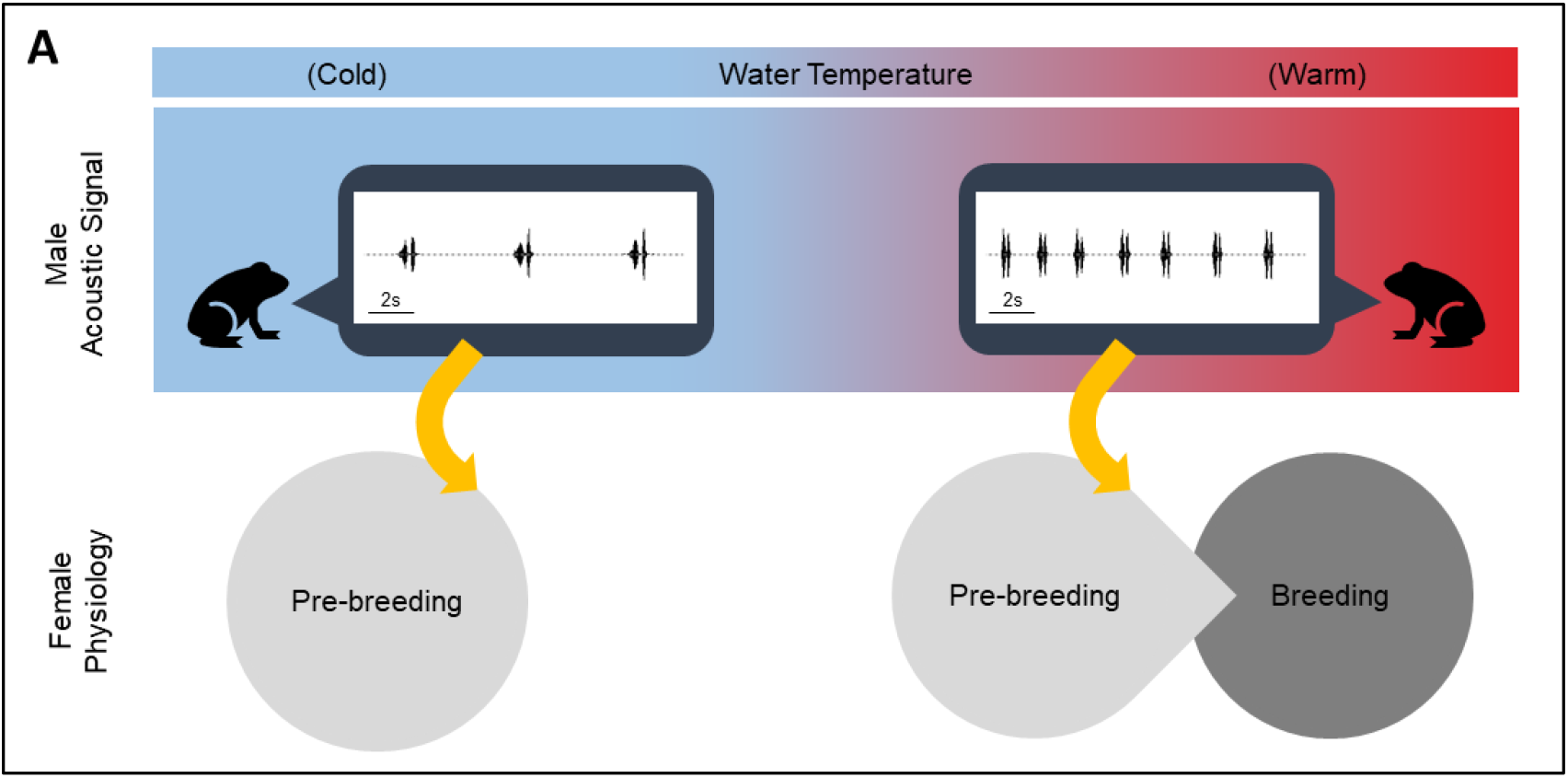
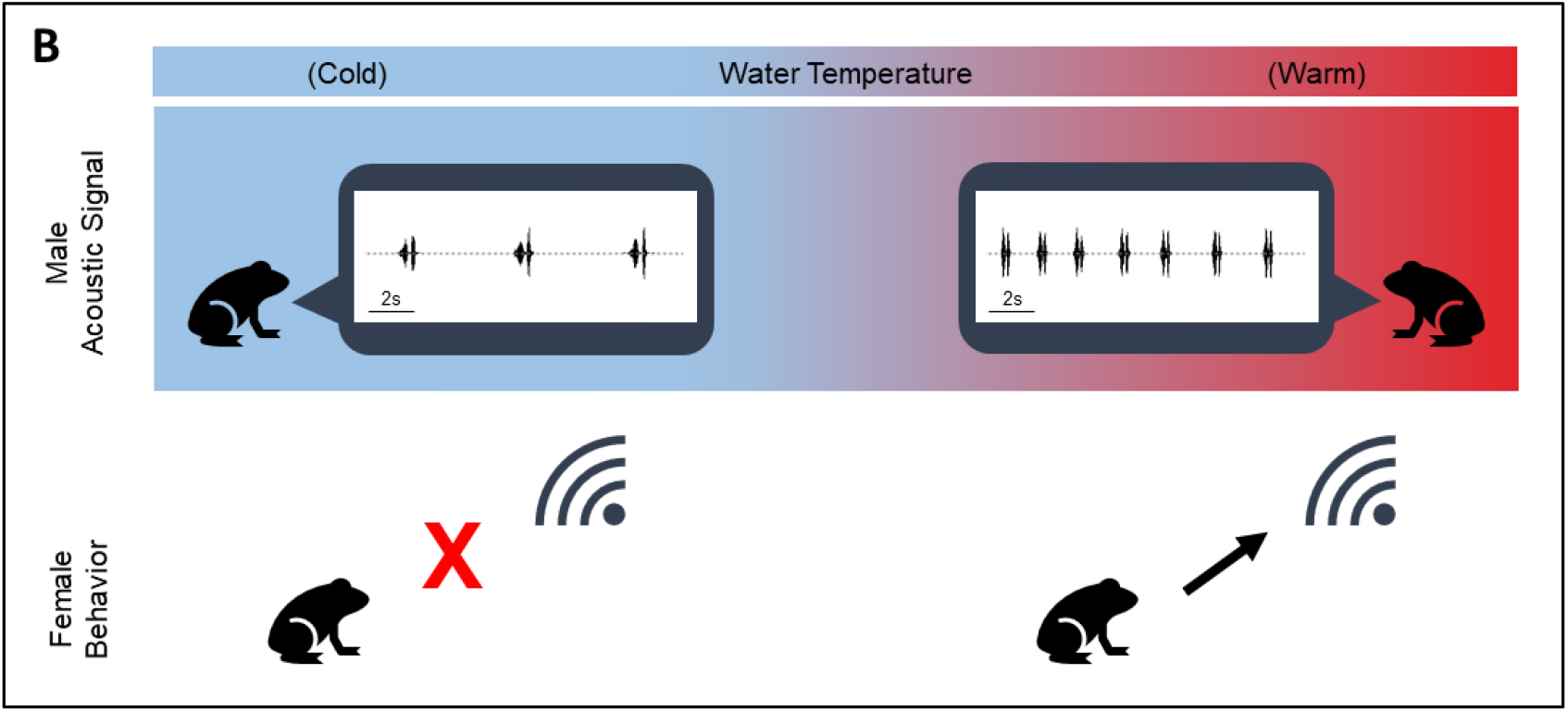
Gross temporal properties of advertisement calls may act as bioclimatic indicators that cue anuran reproduction. A male’s environmental conditions influence the gross temporal properties of his advertisement call. Calls with attractive properties may (A) cue changes in female reproductive physiology and (B) stimulate female reproductive behavior, resulting in navigation to breeding sites and potentially oviposition.

Most anurans follow discrete annual reproductive cycles in which reproductive activity is accompanied by acoustic communication (Rastogi et al., 2011). Over a period of days to months, female anurans undergo follicular development, navigate to breeding sites, approach potential mates using conspecific acoustic signals, and oviposit eggs during amplexus with a male (Wells, 1977, 2010). Due to the essential role of advertisement calls in mediating prezygotic isolation with heterospecifics and mate choice among conspecifics, there may be thresholds or optimal values for the properties of these calls required to cue reproductive processes. Thus, because male advertisement call properties change across the season, reproductive readiness in females may not occur until exposure to amenable environmental conditions, including conspecific advertisement calls with attractive temporal properties (Figure 3A). Similarly, females may only navigate to breeding ponds and approach males that emit advertisement calls with temporal properties above a certain threshold (Figure 3B). While much of phenological research focuses on the minimum thresholds of proximate environmental cues for the onset of a life history event, there may also be maximum thresholds of temporal properties effective for these processes, signaling the end of optimal breeding conditions for the season.

Different behavioral or physiological responses by females to advertisement calls that vary in gross temporal properties should therefore facilitate environmental tracking by constraining annual reproduction to periods in which males emit calls with attractive properties. As increased temperature and advancing seasonality are predicted globally with climate change, our framework predicts that anuran reproduction in systems where seasonality is driven primarily by temperature would correspondingly show advanced dates of annual reproduction. Importantly, in contrast to the more commonly implied mechanisms of organismal responses to proximate environmental cues for reproduction, our framework predicts advancing timing of reproduction with warming as an induced response to a shift in the timing of environmentally informative social signals. In these systems, males may or may not arrive to breeding grounds earlier than historically based on advancing environmental cues for migration, including temperature.

However, because males generally arrive earlier to breeding grounds than females and emit advertisement calls before, during, and after annual oviposition dates, we predict that, regardless of whether male arrival dates advance or not, advertisement calls with gross temporal properties signaling optimal proximate environmental conditions for reproduction would be produced earlier, cueing the organism-level processes contributing to female reproductive behavior and/or physiology earlier (Figure 4). Thus, the gross temporal properties of advertisement calls may function as bioclimatic indicators, facilitating environmental tracking and advancing phenology.

**Figure 4.**
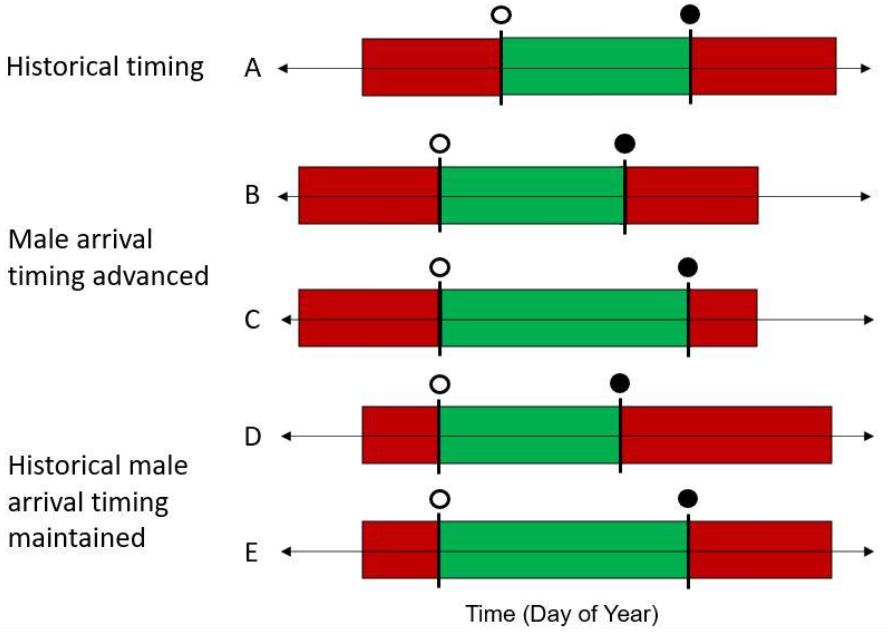
Simplified scenarios of how advancing timing of the production of advertisement calls with attractive gross temporal properties (green shaded bar) may influence the onset (open circle) and cessation (closed circle) of female reproductive activity. Male anurans may produce advertisement calls even at breeding sites with temperatures suboptimal for oviposition, resulting in advertisement calls with unattractive gross temporal properties (red shaded bar), thus the production of attractive calls may advance regardless of whether male arrival timing advances (B, C) or not (D, E). Compared with the historical timing of exposure to advertisement calls with attractive gross temporal properties and reproductive timing (A), advancing the timing of this cue may lead to shifted (B, D) or expanded (C, E) timing of anuran reproductive activity.

Although anuran reproduction is influenced by a complex suite of factors, we highlight two potential organism-driven mechanisms linking a bioclimatic indicator of optimal abiotic conditions to reproduction. To our knowledge, this is the first formalized proposition that seasonally varying conspecific call properties may mediate annual reproductive timing in anurans. The importance of this cue for reproductive physiology and behavior may be species- and population-specific, providing a reliable cue of optimal conditions for breeding in some systems but not others. Detailed discussion of the system-specific factors that may influence the reliability and importance of gross temporal call properties as bioclimatic cues for optimal reproductive conditions, including prolonged versus explosive breeding strategies, temperature-dependent auditory tuning, male calling microhabitat selection, and the presence of heterospecifics, is provided in the supplementary information. Here, we review findings from empirical research demonstrating different physiological and behavioral responses by female anurans in response to male calls that differ in gross temporal properties, then provide guidelines for further research to experimentally test the proposed hypothesis.

### FEMALE RESPONSES TO ADVERTISEMENT CALLS WITH DIFFERENT GROSS TEMPORAL PROPERTIES

We have identified two processes constraining anuran reproduction that may be cued by advertisement calls with threshold values or optimal ranges of gross temporal properties: female reproductive cycling and receptivity to mates. In this section, we briefly review current literature relevant to responses of female anurans to advertisement calls that vary in gross temporal properties. For each process, we highlight areas where further research is needed to understand how these responses may facilitate environmental tracking and influence spatiotemporal patterns of reproduction. The search terms and results used in this literature review are provided in the online supplementary materials.

In social species, changes in female reproductive state may be induced through exposure to male conspecific vocalizations, demonstrated in budgerigars (*Melopsittacus undulatus*; Brockway, 1965), canaries (*Serinus canarius*; Hinde & Steel, 1978), and red deer (*Cervus elaphus*; McComb, 1987). Amphibian reproductive cycling is mediated by the hypothalamic-pituitary-gonadal (HPG) axis, a suite of processes linking proximate cues that stimulate neural activity to the production of neuropeptides in the brain, pituitary hormones, and ultimately gonadal steroids that determines reproductive state (reviewed in Vu & Trudeau, 2016). In anurans, conspecific advertisement calls stimulate neural activity, and, consequently, exposure to conspecific choruses increases the number of immunoreactive gonadotropin-releasing hormone (GnRH-ir) cells at the apex of the HPG axis in males (Burmeister & Wilczynski, 2004).

Exposure to conspecific advertisement calls influences several other processes along the HPG axis, increasing sex steroid production (Burmeister & Wilczynski, 2000; Burmeister & Wilczynski, 2004; Lynch & Wilczynski, 2006) and maintaining gravidity (Brzoska & Obert, 1980; Lea et al., 2001). Despite the integral role of acoustic signals in cueing several physiological processes essential for reproduction, investigation into the effects of variation in conspecific call properties on these processes has typically focused on manipulation of spectral or fine scale temporal properties, such as pulse modulation rate and number of pulses per note (see Brzoska & Obert, 1980; Chu & Wilczynski, 2001; Wilczynski & Ryan, 2010). To our knowledge, no research has yet examined the effects of exposure to acoustic signals that differ in gross temporal properties on any of these physiological processes. Further research is thus needed to determine whether there are threshold or optimal values for the gross temporal properties of conspecific advertisement calls to stimulate physiological processes relevant to reproduction, and whether this relationship facilitates environmental tracking.Gravid female anurans respond behaviorally to conspecific male advertisement calls through navigation to breeding sites, mate choice, and oviposition (Wells, 1977, 2010). However, females show mate selection preferences among calls with different properties (Gerhardt, 1994). The most common gross temporal properties that females select for are call duration and call rate, although preference for intercall interval, call period, and note repetition rate is also evident (see Supplemental Table S3). Because the gross temporal properties of advertisement calls vary with proximate environmental conditions at the breeding site, female preferences for these properties may enable environmental tracking if the preferred properties are characteristic of conditions suitable for oviposition. Further research examining female preference for gross temporal properties as they relate to suitability of conditions at the breeding site is needed to better understand how female behavioral responses may shape spatiotemporal patterns in reproduction.

### RECOMMENDATIONS FOR FUTURE RESEARCH

Despite the well-known role of acoustic communication in anuran reproduction, further research is needed on the physiological and behavioral responses of female anurans to male advertisement calls with gross temporal properties characteristic of conditions that vary in suitability for oviposition. Experimental tests of hypotheses and predictions deriving from the framework proposed here would ideally be designed to distinguish between the effects of proximate abiotic conditions and acoustic signals on female reproductive physiology and behavior. Development of meaningful treatments can be informed through prior field studies of the abiotic conditions associated with reproductive timing and the relationship between temperature and advertisement call properties.

We recommend a factorial experimental design crossing ambient temperature in enclosures with acoustic signals characteristic of male calls under different thermal regimes (Table 1). Acquisition of females outside the breeding season or immediately following oviposition will facilitate collection of longitudinal data on the effects of each experimental treatment on female reproductive state through the event of next oviposition. While the occurrence of, and time to, oviposition are indices of female reproductive behavior, biomarkers of female reproductive physiology can be measured regularly through ovarian monitoring (Calatayud et al., 2018) and hormone assays.

**Table 1.** Example 3×3 factorial design for an experimental test of the proposed hypothesis. Meaningful treatments are chosen based on prior field studies identifying a thermal minimum for reproduction. Three levels for ambient temperature and acoustic environment are crossed for values representative of abiotic conditions and the calls emitted from below (A), at (B), and above (C) the thermal minimum for reproduction (TMR). Dependent variables (female responses) and the intervals at which they are measured will vary based on study constraints, but may include the event of oviposition, number of eggs oviposited, sex steroid concentrations, and the size and number of eggs in the ovary.

|  |  | Acoustic environment |  |  |
| --- | --- | --- | --- | --- |
|  |  | male calls emitted from below TMR | male calls emitted from TMR | male calls emitted from above TMR |
| Ambient temperature | below TMR | female response | female response | female response |
|  | at TMR | female response | female response | female response |
|  | above TMR | female response | female response | female response |

Research resolving the relative contributions of proximate abiotic conditions and acoustic signals on female reproductive physiology and behavior will improve fundamental understanding of the cues constraining anuran reproduction. Consequently, such work can be translated into actionable conservation efforts for species of concern. As amphibians are the most endangered vertebrate class (Wake & Vredenburg, 2008; Barnosky et al., 2011), with an estimated 41% of extant species threatened with extinction (Luedtke et al., 2023), conservation action is urgently needed. First, identifying the cues constraining anuran reproduction will improve predictions of phenological shifts under future climate change, facilitating identification of the most imperiled species and populations. Further, captive breeding programs have been established for many anuran species of concern, but have experienced varied success (Kouba et al., 2009), necessitating the implementation of various assisted reproductive technologies (Silla & Byrne, 2019). Novel approaches informed by this framework, such as exposure of captive breeding populations to playbacks of male advertisement calls with the gross temporal properties identified as most effective at cueing reproduction, may noninvasively improve reproductive success in these populations. Finally, anthropogenic noise can influence the gross temporal properties of calls produced by males and the perception of these properties by females (Zaffaroni-Caorsi et al., 2023). Identification of a strong contribution of acoustic cues to reproduction for a species can inform decision-making for efforts to mitigate anthropogenic noise in areas near breeding sites for species of concern. Broadly, findings from this research may influence several fields of ecology and evolution, spanning advances in fundamental knowledge of anuran reproduction to predictive capability under future climate change and applied conservation management actions.

## CONCLUSION

Through synthesis of literature spanning several fields of ecology and evolution, we have highlighted that:

1. One type of anuran vocalization, the male advertisement call, is fundamental to anuran reproduction, providing a cue for female navigation to breeding sites, reproductive hormone cycling, and mate choice.
2. Gross temporal properties of advertisement calls are dynamic and correlated with the caller’s environment, providing a potential bioclimatic indicator of the suitability of breeding sites for reproduction.
3. Although the effects of variation in gross temporal properties on female anuran reproductive physiology have not been experimentally tested, neural and physiological responses in males to variation in fine scale temporal and spectral properties suggest a potential role for threshold or optimal values of gross temporal properties to likewise cue anuran physiological processes.
4. Anurans can parse variation in gross temporal properties of calls and females show repeatable preferences for gross temporal properties of advertisement calls, with different behavioral responses to calls with different gross temporal properties.

This synthesis proposes that environmentally mediated variation in the gross temporal properties of anuran advertisement calls may act as useful environmental cues for reproduction, with optimal ranges or threshold values for these properties needed to stimulate female physiology and/or behavior. Correspondingly, as gross temporal properties vary with seasonal progression of abiotic conditions or among breeding sites in a heterogeneous landscape, advertisement calls may facilitate environmental tracking and constrain annual reproduction.

Because rising temperatures and advancing seasonality are predicted globally with climate change, these bioindicators may shape spatiotemporal patterns of anuran reproduction. Despite compelling evidence for different physiological and behavioral responses by female anurans to conspecific signals varying in gross temporal properties, experimental tests of the hypotheses and predictions extending from the proposed framework are needed. Findings from this research may improve predictions of anuran phenology under future climate change scenarios and can be translated into actionable conservation management for species of concern.

