## Supplementary Materials for "Anuran call properties as reliable indicators of environmental suitability for reproduction"

*An empirical test of anuran advertisement call properties as reliable indicators of breeding site environmental conditions*

In March 2022, we collected male Sierran tree frogs (*Pseudacris sierra*; N=20) from ephemeral breeding ponds within Quail Ridge Ecological Reserve (Napa County, California, USA) and housed males in outdoor enclosures (10-gallon aquaria with mesh lids; University of California, Davis IACUC protocol #22270). On the night of capture, we measured male mass to the nearest 0.0001g using a digital balance (U.S. Solid) and snout-vent length (SVL) to the nearest 0.01mm using digital calipers (VINCA). Because the dorsal pattern of each *P. sierra* individual is unique and these natural markings can be used to identify individuals, a series of photographs was taken of each frog for individual identification throughout the study (*e.g.*, Bradfield, 2004).

From March 9 through March 11, 2022, we recorded the advertisement calls of individual males in each of three water treatments (chilled, ambient, and warmed) using a repeated measures design. 24 hours prior to the first recording night, males were randomly assigned and exposed to one of the three experimental water treatment, accomplished by placing individual aquaria in communal temperature-controlled water baths. Throughout the 24-hour period, water within each aquarium equilibrated to the water bath temperature. For each subsequent recording night, we again moved aquaria 24 hours prior to recording to a different experimental water bath so that, over the course of the three nights, each frog was recorded in each of the water temperature treatments. Throughout the experiment, the water level within aquaria was consistently maintained at a depth of 4 inches, providing frogs with more than 8 inches of open-air habitat within each aquarium and allowing individuals to exit the water at any time.

However, in all experimental trials, advertisement calls were only emitted by males from within the temperature-controlled water in the tank.

Each night, we recorded each male in a ten-minute trial. Trials were conducted sequentially and randomly, but because males were housed in pairs, we did not allow for males within the same aquarium to be tested consecutively. For each trial, we transferred the non-focal male in the aquarium to a temporary holding container and secured an acoustic recorder (Wildlife Acoustics, Song Meter Mini) and surveillance camera (eufy Security, Outdoor Cam E210) to the lid of the aquarium that still contained the focal male. Following this disturbance, we remotely monitored the focal male from within a nearby structure (approximately 6 meters from the experimental area), obscured from view of the focal frog. After a five-minute acclimation period, we monitored the focal frog over a ten-minute period. If the frog did not call within those ten minutes, the trial was ended and non-occurrence of calling for the focal frog was documented. If the frog did call during the ten-minute period, advertisement calls were acoustically recorded for two minutes starting from the focal frog's first call, after which the trial was ended. Immediately following each trial, we recorded temperature one inch below the water's surface in the aquarium using an automated data logger (Onset, HOBO MX2202), set to record temperature each minute. Due to a power outage that occurred while recording on March 11, 2022, frogs untested on this night were instead tested on March 14, 2022. Males were released to their capture location following the conclusion of the field experiment.

This experiment was repeated in 2023 from March 17 through March 19, with 15 male *P. sierra* collected from the same breeding ponds within Quail Ridge Ecological Reserve. Males were present in large numbers at each site and, through identification using the unique dorsal patterning of each male, we did not identify any frogs recaptured in 2023 from the experiment conducted in 2022. The experimental design remained the same between years, except that, rather than rotating aquaria among water baths, aquaria remained in the same assigned baths across all nights and the experimental treatment imposed on each water bath changed each night. Two equipment changes were made in 2023: we measured water temperature using a digital thermometer with probe (Thermco Products, model ACCD370P) and recorded male calls using a digital field recorder (TASCAM, DR-05X) with microphone attachment (Zoom, ZDM-1 Studio Microphone). In only one out of the total 42 trials in which males called, the focal male emitted aggressive territorial calls away from the water; this male was retested later in the night and, during this trial, emitted advertisement calls from within the temperature-controlled water. Only this second trial was included in data analysis, as our focus is variation in advertisement calls. We measured the gross temporal properties of advertisement calls recorded during experimental trials to the nearest 0.01s using RavenPro acoustic software. We measured call rate (calls per second) for each call group recorded during the 2-minute experimental trial and calculated a weighted average call rate across all call groups per trial. To determine average call duration, we measured and averaged the first 21 consecutive calls per trial.

To determine the relationships among the two gross temporal properties of calls, water temperature within each male's aquarium, and indices of male quality, we created a series of seven candidate models per gross temporal property. We fit each candidate model as a linear mixed effects model (LMM) using maximum likelihood estimation (MLE) with random intercepts for frog ID and date nested within year using the package **lme4** (Bates et al., 2015) in the program R (v4.3.1; R Core Team, 2023). For each response variable (scaled weighted average call rate or scaled average call duration), we first constructed four candidate models, each with only one scaled predictor variable: water temperature or male mass, SVL, or condition (calculated as mass divided by SVL). The three remaining candidate models contained water temperature, one index of male quality, and the interaction between the two predictors. Model diagnostics were assessed through examination of residual plots in the package **DHARMa** (Hartig & Hartig, 2017) and the best fit model for each response variable was determined through comparison of corrected Aikake information criterion (AICc) values (Aikake, 1973; Burnham et al., 2011). The four single-predictor models were then re-run with restricted maximum likelihood estimation (REML) and unscaled variables for estimation and comparison of parameter coefficients and significance (Luke, 2017).

We determined that, under these experimental conditions, call rate and call duration were reliable indicators of water temperature at a male's calling site, regardless of male mass, size, or condition. Through comparison of AICc values for our 7 candidate models (Supplementary Table S1), we identified that, for both call rate and call duration, the model containing only water temperature as a predictor variable was the best fit model. Next, we compared the 95% confidence intervals for each fixed effect estimate from the single-predictor models fit with REML (Supplementary Figure S2) and determined that, for both call rate and duration, only water temperature showed a significant relationship, with no detectable relationships for any measured index of male quality. Finally, from the best fit candidate model containing unscaled water temperature as the only predictor, we determined that water temperature has a significant positive effect on call rate (Supplementary Table S2; $R^2_{marginal} = 0.73$, $\beta = 0.06$, $df = 22.38$, $t = 16.73$, P < 0.001) and significant negative effect on call duration (Supplemental Table S2; $R^2_{marginal} = 0.85$, $\beta = -0.02$, $df = 33.43$, $P < 0.001$). Overall, we found that the gross temporal properties of *P. sierra* advertisement calls are reliable indicators of a male's proximate abiotic environment and, because *P. sierra* males consistently emit advertisement calls from the water in this system, the suitability of the water temperature at breeding sites for reproduction.

*Systematic search for relevant literature*

To identify literature relevant to the framework proposed in this Perspective, we conducted a systematic literature search following established guidelines for ecology and evolution (Foo et al. 2021). The objective of this search was to identify literature relevant to female anuran behavioral and/or physiological responses to advertisement calls with different gross temporal properties. In November 2023, the following terms were searched on Web of Science and analogous terms with engine-specific syntax were searched on Scopus:

TITLE-ABS-KEY ( ( anura* OR frog$ OR toad$ OR treefrog$ ) AND
( breeding* OR reproductive* OR mating* OR advertisement OR social* OR sexual* OR acoustic* OR courtship* ) AND
( call* OR signal* OR cue* OR chorus* OR song* OR vocalization* ) AND
( exposure OR occurrence* OR onset OR presence* OR performance OR temporal OR repetition OR rate$ OR duration OR interval$ OR period$ OR effort OR characteristic* OR propert* ) AND
( female OR intersexual* ) AND
( behavior* OR success OR amplex* OR phonotaxis* OR movement* OR prefer* OR choice* OR choose* OR navigat* OR oviposit* OR physiolog* OR egg* OR follicle* OR vitellogenesis* OR hormon* OR estradiol OR progesterone OR androgen OR testosterone OR GNrH OR neurolog* OR neural* OR inhibit* OR excitat* OR count* ) )

These terms contained exhaustive cross-discipline synonyms specific to our search objectives and resulted in 1,679 unique articles. Each of these articles was manually screened and only included in the review if it met the following criteria: empirical research, including gray literature, on anurans that reported a female physiological or behavioral response to any gross temporal advertisement call property. Because the papers examining physiological responses to gross temporal properties were relatively sparse, our screening criterion for female-only responses was relaxed to include responses by males, as well. In total, only 105 publications met these criteria. To ensure that this list was comprehensive and representative of current literature, we also examined relevant seminal reviews from each of the following fields: animal behavior, ecology, physiology, endocrinology, and neurobiology. As >97% of the relevant references cited within and citing these reviews were identified by our literature search, we believe that the information synthesized here is representative of the literature available on female responses to the gross temporal properties of advertisement calls. While no empirical research examining the influence of gross temporal properties on female anuran reproductive physiology was identified, studies of female behavioral responses to these properties were numerous and spanned several families of anurans (Supplementary Table S3).

*Supplementary discussion*

Due to the diversity of anuran reproductive strategies, the importance of the gross temporal properties of advertisement calls as a cue for reproduction may be species- and population-specific. In some species, primarily those with prolonged breeding seasons, vitellogenesis may begin shortly before oviposition (Mizell, 1964; Jørgensen, 1981). Conversely, in species with explosive breeding seasons, egg maturation can occur prior to overwintering and oviposition occurs soon after emergence in the spring (Iela et al., 1986; Long, 1987; Ritke & Lessman, 1994). In these systems, minimum thresholds of gross temporal properties of advertisement calls for activation of follicular development may not be relevant, although they may still cue receptivity and associated reproductive behaviors. Finally, advancement of exposure to advertisement calls with attractive properties may be limited in explosive breeding systems by advancement of male emergence and arrival at breeding grounds, dependent on length of the period between when males typically arrive at breeding grounds and when oviposition occurs.

Further, species-specific preferences for call properties and auditory tuning may influence the efficacy of environmental tracking via gross temporal properties of advertisement calls in some systems. Sound perception in anurans, as ectotherms, may be influenced by ambient temperature, potentially coupling changes in male call properties with female preferences along a temperature gradient. Temperature-dependent preferences have been demonstrated in phonotactic experiments for pulse repetition rate (Gerhardt, 1978; Gerhardt & Doherty, 1988) and spectral properties (Gerhardt & Mudry, 1980; Humfeld & Grunert, 2015). The neural basis of these preferences for pulse rate have been identified in the sister species *Hyla versicolor* and *H. chrysoscelis*, with thresholds for neural stimulation based on amplitude modulation correlated with ambient temperature in each species (Brenowitz et al., 1985; Rose et al., 1985). However, when manipulated in experimental treatments, variation in call rate and duration influence females' temperature-dependent preferences for pulse rate and, in some cases, even reverse those preferences (Gerhardt & Doherty, 1988). Thus, temperature coupling may occur for only some call properties, with temperature-dependent preferences facilitating species recognition across variable environments while static preferences preserve choices for quality conspecifics.

As discussed throughout this manuscript, the proximate abiotic conditions experienced by males influence the properties of their advertisement calls. Anuran males exhibit diverse calling strategies (reviewed in Wells, 2010): in many species, males call directly from the water at a breeding site and thus the gross temporal properties of advertisement calls serve as reliable indicators of the water temperature at the breeding sites, but males of other species may emit calls from a variety of other microhabitats that vary in distance from the breeding site, including exposed vegetation above the water line, terrestrial sites along the shoreline, and even elevated arboreal perches (Schwartz et al., 2016; De Mello et al., 2018; Chinchilla-Lemus et al., 2020; Martins et al., 2021). Because direct contact may not be made between the water of an aquatic breeding site and these elevated or terrestrial microhabitats, the properties of advertisement calls emitted by males at these calling sites may not be accurate indicators of water temperature within a breeding site. Further, male selection of calling sites may vary with abiotic conditions, with males of some species switching from aquatic to terrestrial calling microhabitats dependent upon temperature (Höbel & Barta, 2014). Although the gross temporal properties of advertisement calls may not directly reflect water temperature of breeding sites in these systems, females may still respond to overall trends in call properties across the breeding season, as the water temperature of shallow depths, where oviposition typically occurs, is strongly correlated with local air temperature across seasons (Livingstone et al., 1999; Winslow et al., 2017).

Finally, population-level factors may also influence the effectiveness of acoustic cues in the proposed framework. Anurans in sympatric populations must weigh the risks of selecting low quality mates against heterospecifics, which results in significantly different selectivity for gross temporal properties in populations where heterospecifics are present versus absent (Gerhardt, 1994; Marquez & Bosch, 1997; Pfennig, 2000; Pfennig & Rice, 2014; Calabrese & Pfennig, 2021).Therefore, sympatric females may be less selective for gross temporal properties than allopatric females, limiting the effect of changes in these properties as a cue for reproduction. Further, abiotic environmental variables that are most limiting to reproduction vary among populations and, depending on climate, may not always include temperature. Although gross temporal properties may be significantly correlated with these optimal conditions, this relationship may not be as robust as the thermal constraints limiting sound production in ectotherms and therefore may not be as reliable a bioclimatic indicator for optimal breeding conditions. Thus, other advertisement call properties may be weighed more heavily in populations that are less constrained by temperature.

### Supplementary Figures

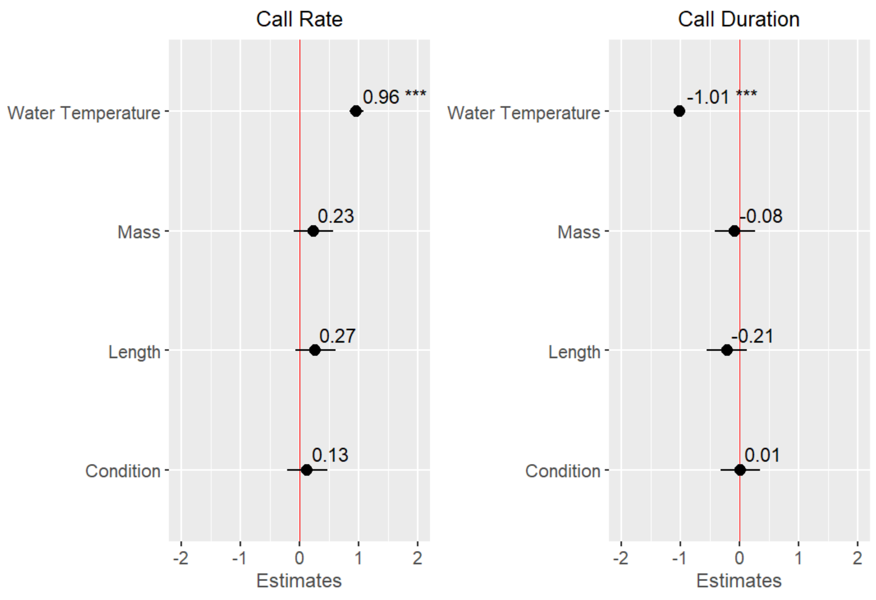


**Supplementary Figure S1. Scaled fixed effects estimates from eight candidate linear mixed effects models predicting call rate (left) or call duration (right) for Sierran chorus frog (*Pseudacris sierra*) advertisement calls in a field experiment conducted at Quail Ridge Ecological Reserve (Napa County, California, USA).** For all eight models, we included random intercepts for frog ID and date, nested within year. Each model contains only one scaled predictor variable: water temperature or the mass, snout-vent length (SVL), or condition (calculated as mass divided by SVL) of the tested frog. Models were fit using restricted maximum likelihood (REML) and the significance of parameters was determined via Satterthwaite approximation, appropriate for small sample sizes (Luke, 2017). Fixed effects estimates are significant (indicated by '***') when the 95% confidence interval does not include a value of 0 (indicated by the red vertical line). We found that, in this system, water temperature is the only predictor variable with a significant effect on weighted average call rate and average call duration.

### Supplementary Tables

**Supplementary Table S1.** Differences in corrected Aikake information criterion (AICc) values (Aikake, 1973; Burnham et al., 2011) among candidate models predicting weighted average call rate or average duration in a field experiment determining the effects of water temperature and male quality on *Pseudacris sierra* advertisement call gross temporal properties. The best fit model for each response variable is indicated by a bolded value of 0.

| Predictor variable(s) \ Response variable | Call rate | Call duration |
|---|---|---|
| Water temperature | **0** | **0** |
| Mass | 65.35 | 94.64 |
| SVL | 64.86 | 93.28 |
| Condition | 66.76 | 94.88 |
| Water temperature * Mass | 5.62 | 4.39 |
| Water temperature * SVL | 5.50 | 5.85 |
| Water temperature * Condition | 5.87 | 4.02 |

**Supplementary Table S2.** Output from two linear mixed models (LMMs), fit through restricted maximum likelihood (REML), with random intercepts for frog ID and date, nested within year, and water temperature as the predictor variable. The response variable of the first model is weighted average call rate and, for the second, average call duration. We determined the significance of parameter estimates via Satterthwaite approximation, appropriate for small sample sizes (Luke, 2017).

| | **Call Rate (calls/s)** | | | **Call Duration (s)** | | |
|---|---|---|---|---|---|---|
| *Predictors* | *Estimates* | *CI* | *p* | *Estimates* | *CI* | *p* |
| (Intercept) | 0.01 | -0.10 – 0.13 | 0.842 | 0.51 | 0.48 – 0.54 | **<0.001** |
| Water Temperature (C) | 0.06 | 0.05 – 0.06 | **<0.001** | -0.02 | -0.02 – -0.01 | **<0.001** |
| **Random Effects** | | | | | | |
| $\sigma^2$ | 0.01 | | | 0.00 | | |
| $\tau_{00}$ | $0.02_{\text{FrogID}}$ | | | $0.00_{\text{FrogID}}$ | | |
| | $0.00_{\text{Date:Year}}$ | | | $0.00_{\text{Date:Year}}$ | | |
| | $0.00_{\text{Year}}$ | | | $0.00_{\text{Year}}$ | | |
| ICC | 0.74 | | | 0.56 | | |
| N | $7_{\text{Date}}$ | | | $7_{\text{Date}}$ | | |
| | $2_{\text{Year}}$ | | | $2_{\text{Year}}$ | | |
| | $23_{\text{FrogID}}$ | | | $23_{\text{FrogID}}$ | | |
| Observations | 42 | | | 42 | | |
| Marginal $R^2$ / Conditional $R^2$ | 0.733 / 0.931 | | | 0.849 / 0.933 | | |

**Supplementary Table 3.** Empirical studies in which at least one gross temporal property of conspecific male advertisement calls was significantly related to an index of anuran female preference. Studies are organized alphabetically by taxonomic family.

| Species | Significant gross temporal property | Response measured | References |
|---|---|---|---|
| **Alytidae** | | | |
| *Alytes cisternasii* | call duration | phonotaxis | Marquez & Bosch (1997) |
| | call rate | phonotaxis | Bosch & Márquez (1996); Márquez et al. (2008) |
| | | reciprocal female call | Bosch (2001) |
| | intercall interval | phonotaxis | Bosch & Márquez (2005) |
| *Alytes muletensis* | call rate | phonotaxis | Dyson et al. (1998) |
| *Alytes obstetricans* | call duration | phonotaxis | Marquez & Bosch (1997) |
| | | male mating success | Lodé & Jacques (2003) |
| | call rate | phonotaxis | Bosch & Márquez (1996); Márquez et al. (2008) |
| **Aromobatidae** | | | |
| *Allobates femoralis* | call duration | mate choice | Peignier et al. (2022) |
| *Anomaloglossus [Colostethus] beebei* | call duration | Phonotaxis | Pettitt et al. (2020) |
| | call rate | phonotaxis | Bourne et al. (2001) |
| **Bufonidae** | | | |
| *Anaxyrus [Bufo] americanus* | call duration | Phonotaxis | Sullivan (1992) |
| | call rate | Phonotaxis | Sullivan (1992) |
| *Anaxyrus [Bufo] cognatus* | call duration | Phonotaxis | Leary et al. (2006) |
| *Anaxyrus [Bufo] woodhousei* | call rate | Phonotaxis | Sullivan (1982); Sullivan (1983) |
| | | male mating success | Sullivan (1982); Sullivan (1983); Sullivan (1989) |
| *Bufotes [Bufo] viridis* | call duration | Phonotaxis | Castellano et al. (2000) |
| | Intercall interval | Phonotaxis | Castellano & Giacoma (1998) |
| *Epidalea [Bufo] calamitas* | call rate | Phonotaxis | Arak (1998) |
| *Incilius [Bufo] valliceps* | call duration | Phonotaxis | Wagner & Sullivan (1995) |
| | call rate | Phonotaxis | Wagner & Sullivan (1995) |
| | | male mating success | Wagner & Sullivan (1995) |
| *Sclerophrys capensis [=Bufo rangeri]* | call rate | Phonotaxis | Cherry (1993) |
| **Dendrobatidae** | | | |
| *Dendrobates leucomelas* | call duration | male mating success | Forsman & Hagman (2006) |
| | call rate | male mating success | Forsman & Hagman (2006) |
| *Epipedobates tricolor* | call duration | male mating success | Forsman & Hagman (2006) |
| | call rate | male mating success | Forsman & Hagman (2006) |
| *Hyloxalus [Colostethus, Dendrobates] subpunctatus* | note repetition rate | Phonotaxis | Lüddecke (2002) |
| *Oophaga [Dendrobates] pumilio* | call rate | Phonotaxis | Pröhl (2003) |
| **Dicroglossidae** | | | |
| *Quasipaa spinosa* | call duration | Phonotaxis | Yu et al. (2020) |
| | call rate | phonotaxis | Yu et al. (2020) |
| **Eleutherodactylidae** | | | |
| *Eleutherodactylus coqui* | call rate | Phonotaxis | Lopez & Narins (1991) |
| | | mate choice | Lopez & Narins (1991) |
| **Hylidae** | | | |
| *Agalychnis moreletii* | call duration | male mating success | Briggs (2010) |
| | intercall interval | male mating success | Briggs (2010) |
| *Dendropsophus bipunctatus* | call rate | Phonotaxis | Wogel & Pombal (2007) |
| *Dendropsophus carnifex* | call rate | Phonotaxis | Oliva et al. (2010) |
| *Dendropsophus [Hyla] microcephala* | call rate | Phonotaxis | Schwartz (1986) |
| *Dryophytes [Hyla] avivoca* | call duration | Phonotaxis | Martínez-Rivera & Gerhardt (2008) |
| *Dryophytes [Hyla] chrysoscelis* | call duration | Phonotaxis | Vélez et al. (2013); Tanner et al. (2017) |
| | | mate choice | Morris & Yoon (1989) |
| | | inertial energy | Gupta & Bee (2023) |
| | call rate | Phonotaxis | Ward et al. (2015); Tanner et al. (2017) |
| | | mate choice | Morris & Yoon (1989) |
| *Dryophytes [Hyla] cinerea* | call duration | Phonotaxis | Gerhardt (1987); Höbel (2010) |

| | | | |
|---|---|---|---|
| | call rate | Phonotaxis | Gerhardt (1987); Humfeld (2008); Höbel (2010); Davis & Leary (2015); Laird et al. (2016) |
| *Dryophytes [Hyla] gratiosa* | call duration | Phonotaxis | Burke & Murphy (2007); Poole & Murphy (2007) |
| | call rate | Phonotaxis | Murphy & Gerhardt (1996); Murphy & Gerhardt (2000); Burke & Murphy (2007) |
| *Dryophytes [Hyla] squirella* | call rate | Phonotaxis | Taylor et al. (2007) |
| *Dryophytes [Hyla] versicolor* | call duration | phonotaxis | Klump & Gerhardt (1987); Sullivan & Hinshaw (1992); Gerhardt (1994); Gerhardt et al. (2000); Schwartz et al. (2001); Gordon & Gerhardt (2009); Kolodziej (2014); Reichert & Höbel (2015); Underhill & Höbel (2017); Stratman & Höbel (2019); Feagles & Höbel (2022) |
| | | inertial energy | Gupta & Bee (2023) |
| | call rate | phonotaxis | Sullivan & Hinshaw (1992); Gerhardt et al. (1996); Gordon & Gerhardt (2009) |
| | intercall interval | phonotaxis | Boyd & Gordon (2021) |
| | call period | phonotaxis | Reichert & Höbel (2015); Stratman & Höbel (2019) |
| *Hyla arborea* | call duration | phonotaxis | Richardson et al. (2010) |
| | call rate | phonotaxis | Richardson et al. (2010) |
| *Hyla intermedia* | call duration | phonotaxis | Botto & Castellano (2016) |
| | | male mating success | Castellano et al. (2009) |
| | call rate | phonotaxis | Castellano (2010); Botto & Castellano (2016) |
| | | male mating success | Castellano et al. (2009) |
| *Pseudacris crucifer* | call duration | phonotaxis | Doherty & Gerhardt (1984) |
| *Pseudacris triseriata* | call duration | phonotaxis | Martof & Thompson (1964) |
| | call rate | phonotaxis | Martof & Thompson (1964) |
| **Hyperoliidae** | | | |
| *Hyperolius marmoratus* | call duration | phonotaxis | Grafe (1997) |
| | call rate | phonotaxis | Passmore et al. (1992); Grafe (1997) |
| | | male mating success | Passmore et al. (1992); Grafe (1997) |
| **Leptodactylidae** | | | |
| *Engystomops pustulosus* | call duration | phonotaxis | Dawson & Ryan (2009) |
| | intercall interval | phonotaxis | Bosch et al. (2000) |
| *Physalaemus fischeri* | call duration | phonotaxis | Tárano & Herrera (2003) |
| *[=enesefae]* | intercall interval | phonotaxis | Tárano & Herrera (2003) |
| **Myobatrachidae** | | | |
| *Crinia georgiana* | call rate | phonotaxis | Smith & Roberts (2003) |
| | | male mating success | Smith & Roberts (2003) |
| *Pseudophryne corroboree* | call duration | male mating success | Kelleher et al. (2022) |
| | call rate | male mating success | Kelleher et al. (2022) |
| **Ranidae** | | | |
| *Babina daunchina* | call duration | phonotaxis | Cui et al. (2012) |
| **Scaphiopodidae** | | | |
| *Scaphiopus couchii* | call duration | phonotaxis | Pfennig & Tinsley (2002) |
| *Spea multiplicata* | call rate | phonotaxis | Pfennig (2000); Pfennig & Rice (2014); Calabrese & Pfennig (2021); Calabrese & Pfennig (2022) |
